# Differential expression of microRNAs in Alzheimer’s disease brain, blood and cerebrospinal fluid: a systematic review and meta-analysis

**DOI:** 10.1101/499491

**Authors:** Petros Takousis, Angélique Sadlon, Jessica Schulz, Inken Wohlers, Valerija Dobricic, Lefkos Middleton, Christina M. Lill, Robert Perneczky, Lars Bertram

## Abstract

**INTRODUCTION:** Several microRNAs (miRNAs) have been implicated in Alzheimer’s disease (AD) pathogenesis but the evidence from individual case-control-studies remains inconclusive.

**METHODS:** A systematic literature review was performed, followed by standardised multi-stage data extraction, quality control, and meta-analyses on eligible data for brain, blood, and cerebrospinal fluid (CSF) specimens. Results were compared with miRNAs reported in the abstracts of eligible studies or recent qualitative reviews to assess novelty.

**RESULTS:** Data from 147 independent datasets across 107 publications were quantitatively assessed in 461 meta-analyses. Twenty-five, five, and 32 miRNAs showed study-wide significant differential expression (α<1.08×10^−4^) in brain, CSF, and blood-derived specimens, respectively, with 5 miRNAs showing differential expression in both brain and blood. Of these 57 miRNAs, 13 had not been reported in the abstracts of previous original or review articles.

**DISCUSSION:** Our systematic assessment of differential miRNA expression is the first of its kind in AD and highlights miRNAs of relevance.

## 1. BACKGROUND

In addition to genetic variants, epigenetic factors may play an important role in AD pathogenesis, such as DNA methylation or the regulatory effects of small non-coding RNAs, in particular microRNAs (miRNAs). MiRNAs are 18-25 nucleotides in length and primarily regulate gene expression at the post-transcriptional level, through recognition of specific binding sites located mainly in the 3’-untranslated region of their target messenger RNAs (mRNAs) [1]. Thus, changes in miRNA expression can lead to translational repression and, consequently, reductions in the levels of the respective proteins.

Due to their crucial role in “fine-tuning” gene expression, miRNAs have been proposed as biomarkers and/or therapeutic targets for a range of human disorders, including AD [2–4]. A growing body of literature suggests that miRNAs are causally linked to AD by directly affecting the underlying pathogenic pathways, e.g. by targeting *APP* [5] or *BACE1* expression [6, 7], thereby altering the risk and/or progression of the disease [4, 5]. However, despite their possible pathophysiological relevance in AD and their potential to serve as biomarkers, there is still no conclusive picture as to which miRNAs play the lead roles, e.g. by showing consistent patterns of differential expression in AD vs healthy controls. Furthermore, interpretation of the existing data is severely hampered by methodological shortcomings of the individual studies, including small sample size and methodological heterogeneity, making it difficult to compare different studies.

The present work aimed at performing a systematic review and meta-analysis of all published studies probing for differential expression of miRNAs in AD cases vs controls in brain, blood, and cerebrospinal fluid (CSF) specimens. The main goal was to identify miRNAs which show significant and consistent differential expression when comparing results across publications. In line with similar efforts applied to results from genetic association studies developed by members of our group [8, 9], we applied a standardised multi-stage data extraction and quality control protocol followed by meta-analysis on all resulting eligible data. Our work, which represents the first of its kind in AD, highlights 57 unique miRNAs showing consistent and highly significant differential expression across studies. This systematic review generates a knowledge-base which could prove instrumental in future studies assessing the role of miRNAs in AD pathogenesis and their potential as AD biomarkers.

## 2. METHODS

See Figure 1 for a quantitative summary of the various steps underlying this systematic review; in addition, the Supplementary Material provides a full description of all methods applied in this study. In the following section, methods are only summarised to highlight the most essential features and check-points of our study. Overall, the work-flow and data collection procedures applied in this work are similar to those for genetic association studies developed previously by members of our group [8, 9]. The approach was adapted here to the characteristics of gene expression studies. In brief, this entailed a systematic literature search for miRNA expression studies in AD using the PubMed database, the systematic extraction of data from eligible studies into a project-specific database, and a range of data cleaning and reformatting steps followed by extensive plausibility and quality checking (e.g. alignment of miRNA identifiers to those listed in miRbase, v21 (http://www.mirbase.org), identification of duplicate publications and/or sample overlap across studies, and double checking of entries by independent team members). Statistical analyses of the remaining quality controlled data entailed meta-analysing study-level results for miRNAs with at least three independent data points per specimen.

**Figure 1.**
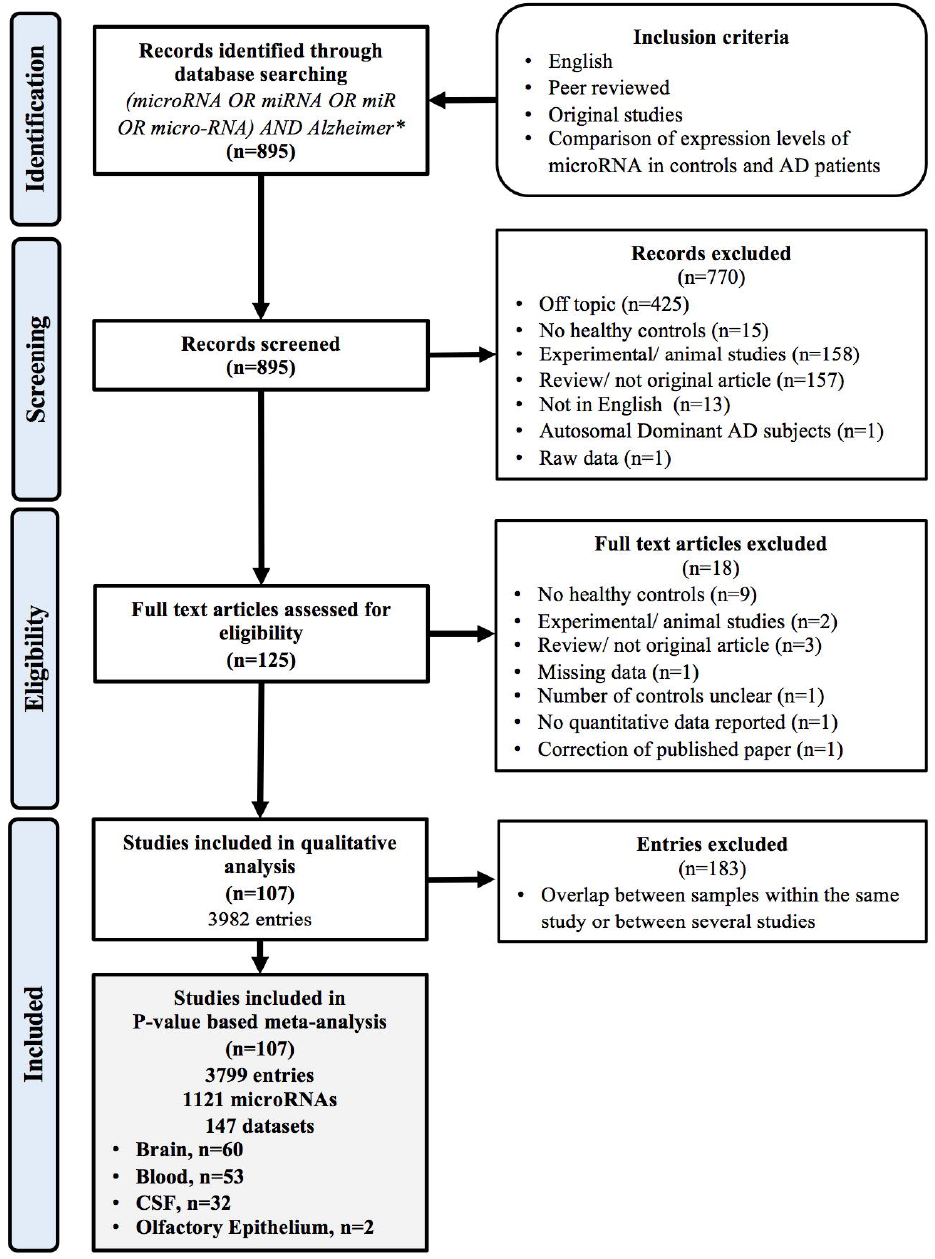
Work-flow implemented to identify eligible datasets

The meta-analyses themselves were performed using a customised R script (https://www.r-project.org; available upon request) applying Stouffer’s method,[10] i.e. converting reported *p*-values into signed z-scores, which were then meta-analysed using inverse variance weighting based on sample size, a method very similar to that described previously for the meta-analysis of GWAS [11]. Thresholds of statistical significance were defined after Bonferroni correction for multiple testing. This was based on the overall number of meta-analyses (n=461) performed across all three strata resulting in a study-wide *α* of 0.05/461=1·08×10^−4^. Target genes of miRNAs showing study-wide significance were identified and evaluated based on next-generation sequencing data from human post-mortem brain samples [12] and experimentally validated miRNA-target interactions from miRTarBase [13]. Finally, we performed genetic enrichment analyses using these target gene predictions based on GWAS summary statistics from the International Genomics of AD Project (IGAP) [14].

## 3. RESULTS

### 3.1 Study characteristics

Our systematic literature search yielded a total of 895 publications, of which 107 were classified as eligible for inclusion after careful, multi-stage screening (Figure 1). After quality control, data from 147 independent datasets across these 107 publications were subsequently extracted, entered into our database and included in the meta-analyses. Possible reasons for exclusion of publications/datasets from meta-analysis are summarised in Figure 1.

MiRNA expression data included in the meta-analyses were generated from specimens classified as derived from “brain”, “CSF”, and “blood”. Of all 147 meta-analysed datasets, 60 were generated from brain, 32 from CSF, and 53 from blood specimens. Two additional datasets utilised miRNA expression data from anterior nasal septum mucosa and olfactory bulb (Supplementary Table 1). Fifteen publications reported results from more than one specimen type: two from brain and CSF [5, 15], four from brain and blood [16–19], and nine from CSF and blood [20–28]. Datasets included in the brain meta-analyses comprised specimen from different regions: basal forebrain/entorhinal (n=3), hippocampus/medial temporal lobe (n=31), frontal/prefrontal lobe (n=14) and parietal lobe (n=2). For four datasets, brain regions were not specified; six datasets consisted of two or more brain regions (Supplementary Table 1). The median sample size per dataset was 34·5 across all studies (interquartile range [IQR] 17-68), 19 for brain (IQR 12-34), 38 for CSF (IQR 21-62), and 68 for blood (IQR 37-97).

Overall, data for a total of 1,121 different miRNAs were reported across all studies. The median number of meta-analysed miRNAs analysed per study was four (IQR 2-13). Only nine studies used a large-scale approach, defined here as reporting findings on more than 100 miRNAs, and data from all of these studies were included in our meta-analyses (Supplementary Table 1). Of all included miRNAs, 295 had been assessed in at least three independent datasets in at least one specimen type and were, thus, deemed eligible for metaanalysis. One hundred and twenty-three of the 295 miRNAs were meta-analysed in more than one specimen type, i.e. 58 in brain and blood, 20 in brain and CSF, two in CSF and blood, and 43 in all three specimen types. Overall, this resulted in sufficient data to conduct 461 individual meta-analyses for the present study (Supplementary Table 2).

### 3.2 Meta-analysis results

Two hundred and sixty meta-analyses were based on data collected using brain-, 66 using CSF-, and 135 using blood-derived specimens. The median number of datasets included per meta-analysis across all miRNAs was four in brain and blood, and three in CSF, with maximum values of 19, 10, and 11, respectively. The median combined sample size across all miRNAs in brain, CSF, and blood was 42·5 (IQR 23-85), 164 (IQR 98-205), and 259 (IQR 195-318), respectively.

Twenty-five of the 260 (10%) miRNAs meta-analysed for brain showed study-wide significant differential expression in AD cases vs controls with *p*-values ranging from 4·13×10^−13^ to 9·10×10^−5^ (Table 1). The three miRNAs with the most statistically significant differential expression were hsa-miR-125b-5p, hsa-miR-501-3p, and hsa-miR-885-3p. Twelve miRNAs were up-regulated, while 13 were downregulated in AD compared to controls. In addition, 97 brain miRNAs showed nominally (*α*=0·05), but not study-wide, significant differential expression (Supplementary Table 2). In CSF, five of the 66 (8%) meta-analysed miRNAs showed study-wide significant differential expression in AD cases vs controls with *p*-values ranging from 3.48×10^−7^ to 3.76×10^−5^ (Table 1). All five were downregulated in AD compared to controls. The three miRNAs with the most statistically significant differential expression in CSF were hsa-miR-598-3p, hsa-miR-451a, and hsa-miR-9-5p. In addition, 23 CSF miRNAs showed nominally significant differential expression (Supplementary Table 2). Finally, in blood, thirty-two of 135 (24%) meta-analysed miRNAs showed study-wide significant differential expression in AD cases vs controls with *p*-values ranging from 916×10^−24^ to 4·64×10^−5^ (Table 1). Thirteen miRNAs were upregulated, while 19 were downregulated in AD cases vs controls. The three miRNAs with the most statistically significant differential expression were hsa-miR-342-3p, hsa-miR-191-5p, and hsa-let-7d-5p. Thirty-eight additional miRNAs showed nominally significant differential expression (Supplementary Table 2).

**Table 1.**
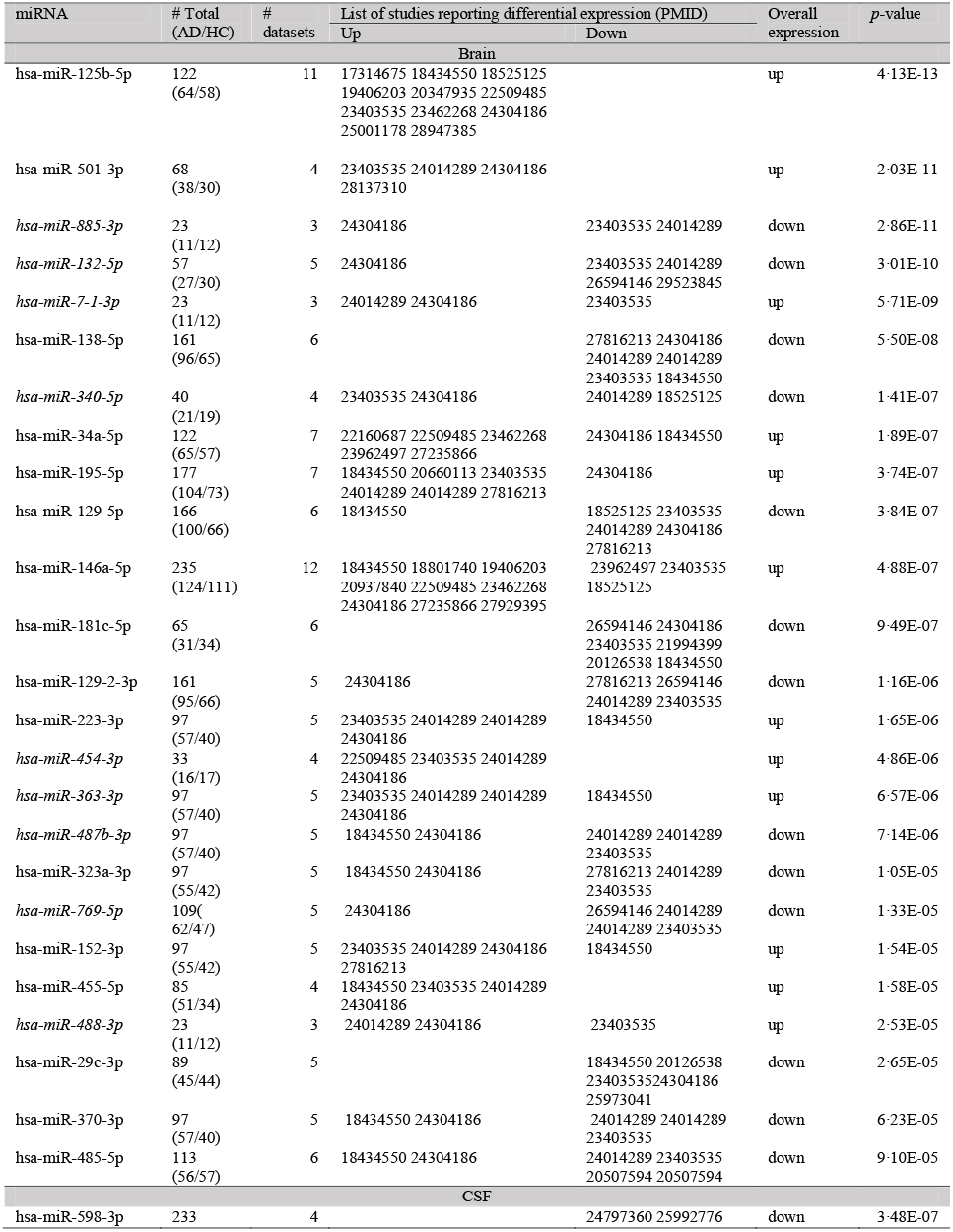
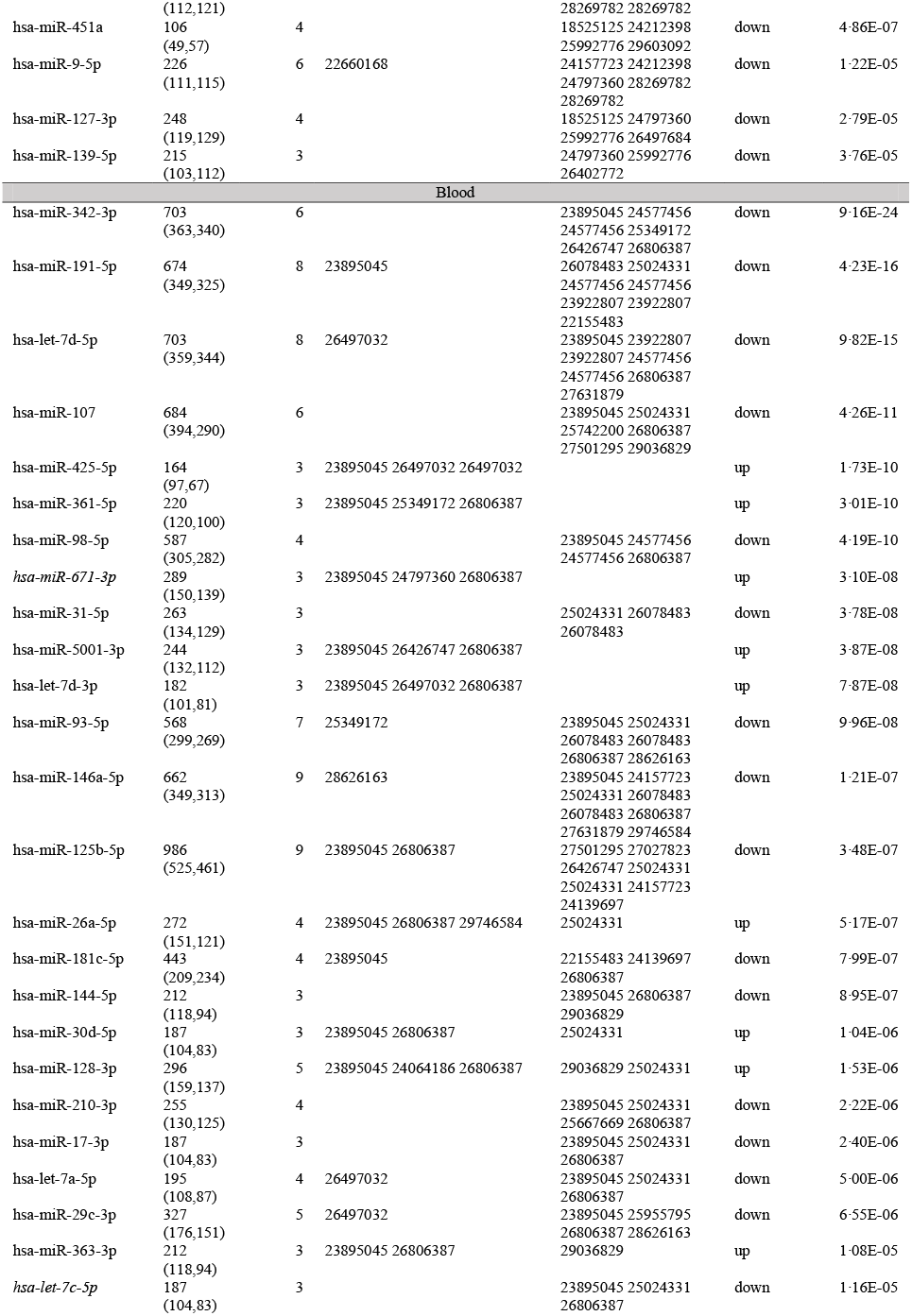
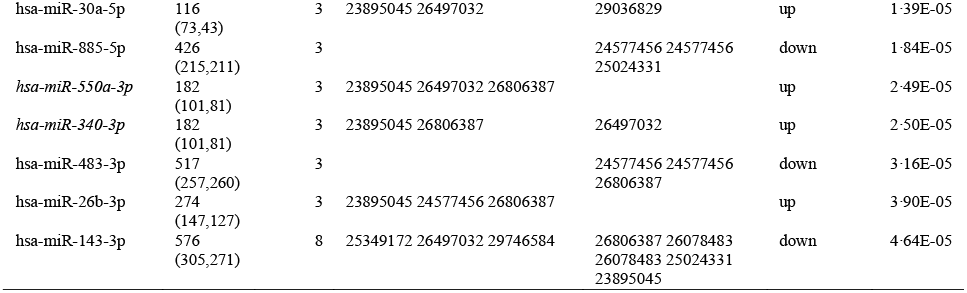
Significant meta-analysis results of differentially expressed miRNAs in brain, CSF and blood in Alzheimer’s disease patients and controls (miRNAs not reported in the abstracts of previous original articles or in qualitative reviews, in italics).

Interestingly, five miRNAs showed study-wide significant differential expression in both brain and blood, i.e. hsa-miR-181c-5p and hsa-miR-29c-3p were downregulated in both, whereas hsa-miR-125b-5p, hsa-miR-146a-5p, and hsa-miR-223-3p were upregulated in brain and downregulated in blood (Table 1). No overlaps in up-or down-regulated miRNAs showing study-wide significant differential expression were observed between CSF and either brain or blood.

### 3.3 MiRNA target identification and genetic enrichment analyses

To map miRNA-target interactions of the study-wide significant miRNAs we used two sources of information. First, we interrogated Ago2 HITS-CLIP sequencing results derived from human brain samples [12], which highlighted nine genes previously implicated in AD (see Supplementary Material on how these were defined), i.e. *<u>APP, CCDC6, CD2AP, CLU</u>, CNTNAP2, <u>FERMT2, PICALM</u>, PTK2B*, and *SORL1* (Figure 2 and Supplementary Table 3a). Second, we used experimentally validated miRNA-target interactions posted on miRTarBase [13], which revealed 12 known AD genes, i.e. *ADAMTS4, <u>APP, CCDC6, CD2AP, CLU, FERMT2</u>, HS3ST1, <u>PICALM</u>, PLCG2, SCIMP, SLC24A4*, and *SPI1* (Supplementary Table 3b; overlapping targets across both approaches are underlined). These observations provide additional support for a genuine involvement of the implicated miRNAs in AD pathogenesis. In addition to these known AD genes, each miRNA was predicted to also interact with several targets not previously linked to AD. We were, thus, interested to assess whether the full sets of miRNA-specific targets showed an enrichment (using PASCAL) [29] of significant GWAS signals based on genome-wide summary statistics published along with the IGAP study [14]. While three nominally significant enrichments were observed, i.e. with hsa-miR-17-3p, hsa-miR-7-1-3p, and hsa-miR-93-5p (Supplementary Table 3), none of these results reached study-wide significance using false discovery rate (FDR) controlling for multiple comparisons. Future work needs to assess the potential functional role, if any, of these non-AD genes targeted by the miRNAs highlighted in our meta-analyses.

**Figure 2.**
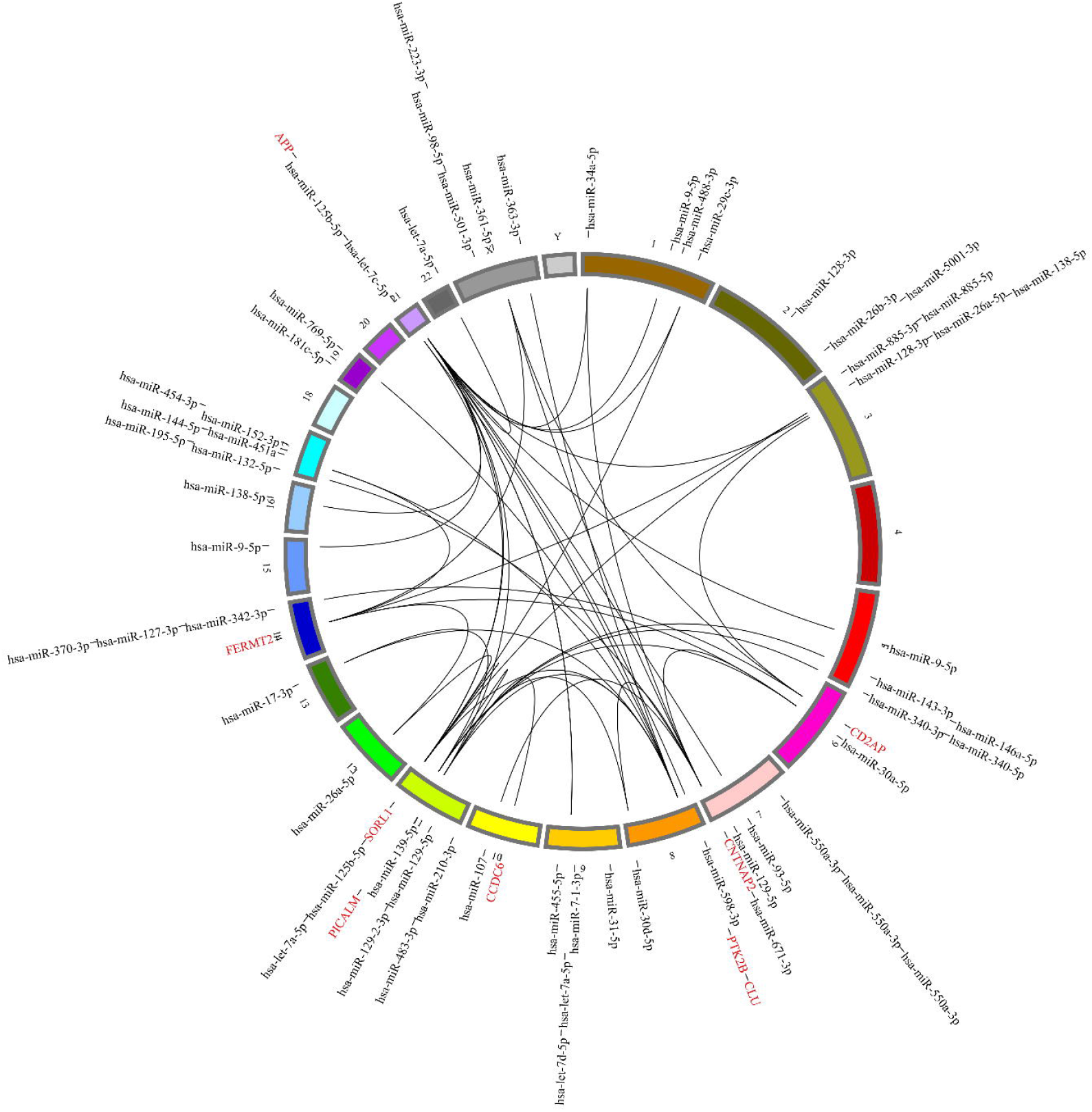
Circos plot of chromosomal locations of all miRNAs showing study-wide significant differential expression in the meta-analyses of AD cases vs controls. Connecting lines are between miRNAs and their target genes (as determined by Ago2 HITS-CLIP sequencing in post-mortem human brain)[12] showing genome-wide significant associations with AD risk from recent GWAS (see Supplementary Material). For more details on the AD-specific miRNA-target pairs displayed in this figure see Supplementary Table 3A.

### 3.4 Comparison of meta-analysis results to qualitative summaries of differential miRNA expression in Alzheimer’s disease

We generated two sets of qualitative reference data. First, we recorded miRNAs highlighted in the abstracts of the 107 publications forming the literature knowledge base of our study and compared these miRNAs to our meta-analysis results. In total, there were 124 different unique miRNAs reported in the abstracts of articles included in our study at varying levels of frequency (Supplementary Figure 1). Thirty-one (25%) of these miRNAs were also found to be differentially expressed at study-wide significance, and another 40 (i.e. overall 57%) at nominal significance in our meta-analyses (Supplementary Figure 1). For the remaining 53 miRNAs either no significant differential expression was observed here (i.e. for 30 [24%] miRNAs), or meta-analyses could not be performed due to a lack of sufficient data (i.e. for 23 [19%] miRNAs). Most importantly, 26 miRNAs that were found to be differentially expressed at study-wide significance here had not been reported in the abstract of any of the individual research articles.

Second, we compared our quantitative findings with miRNAs highlighted in 17 recent qualitative reviews published since 2016 (see Supplementary Material for details on how publications were selected). Overall, these articles reported 201 miRNAs as being potentially relevant in AD pathogenesis but only 40 (20%) of these miRNAs showed study-wide significant differential expression in our meta-analyses. Also counting nominally significant results extended this list to another 62 (i.e. overall 51%) overlapping miRNAs. Of the remaining 99 miRNAs, 49 (24%) showed no evidence of differential expression here, while 50 (25%) did not have sufficient data to be meta-analysed. A total of 17 miRNAs showing study-wide significant differential expression in our analyses were not highlighted in any of the 17 qualitative review articles. Four of these overlapped with miRNAs reported in abstracts of original articles (see above), leaving a total of 13 miRNAs (out of 57, i.e. 23%) which can be considered “novel” results of this study (italicised miRNA names in Table 1).

## 4. DISCUSSION

This work represents a comprehensive and quantitative account of studies assessing differential miRNA expression in AD cases vs controls across three specimen types. We utilised a multi-pronged protocol combining systematic literature searches with rigorous data extraction, quality control and double-checking procedures, followed by meta-analysis of eligible study data. The approach applied here was originally developed for and successfully applied to genetic association studies in AD and other fields [8, 9], adapted to the specifics of studying miRNA expression. To the best of our knowledge, our study represents the first comprehensive systematic synthesis and quantitative assessment of miRNA differential expression data in the field of AD research.

Taken together, we identified 57 unique miRNAs showing study-wide significant (i.e. P<1.08×10^−4^) differential expression in brain-, CSF-, and/or blood-derived specimens. Five of these miRNAs showed evidence for study-wide differential expression in more than one tissue type and several of these miRNAs target established AD genes which may suggest their involvement in AD pathogenesis. Forty-four of the differentially expressed miRNAs were highlighted in the abstracts of publications eligible for data extraction in this study and/or qualitative review papers on the role of miRNA expression in AD. This means that nearly one quarter (i.e. 13) of the miRNAs identified here were not highlighted in previous publications and can, thus, be considered “novel” findings of our systematic analyses. Notably, nine of these 13 novel miRNAs were found to be differentially expressed in brain-derived specimen and according to published functional data, many of the novel miRNAs identified may play important roles in AD-related mechanisms (summarised in Supplementary Table 4). Alongside this publication we make the complete knowledge base collated for this study available to the community (Supplementary Table 5): this includes the database of all study-level miRNA expression data (with nearly 4,000 entries across all 107 included studies) and details of all 461 meta-analysis results (in Supplementary Table 2).

More than half of the miRNAs showing study-wide significant differential expression (i.e. 32 of 57) were identified by meta-analyses performed on blood-derived specimens. This finding could prove an important prerequisite for future investigations aimed at identifying miRNA signatures able to differentiate between healthy aging and AD, potentially in the prodromal stage. Further research into this topic is required to validate and extend our findings in prospective studies, e.g. to assess the diagnostic and prognostic properties of the miRNAs highlighted by meta-analysis here. Such blood-based miRNA signature(s), perhaps in concert with other blood-based biomarkers and genetic predictors, would facilitate risk stratification for large-scale prevention programs and provide a window of opportunity for early therapeutic intervention. Such signature(s) could also serve as a tool to assess disease stage and monitor disease progression or therapeutic effects.

It is particularly noteworthy that five miRNAs showed study-wide significant differential expression in both brain and blood: hsa-miR-181c-5p and hsa-miR-29c-3p were found to be downregulated in both, whereas hsa-miR-125b-5p, hsa-miR-146a-5p, and hsa-miR-223-3p were upregulated in brain and downregulated in blood (Table 1). The fact that the same miRNAs show differential expression in different specimen types in AD vs controls emphasises their potential role in AD and highlights their potential as future biomarkers. Different explanations have been proposed to explain the observation of opposing effect directions. For example, if the pathology is associated with a tissue-specific concentration change of a ubiquitously-expressed miRNA, the effect on circulating miRNA concentration could be minimal since only a fraction of it derives from the affected tissue [30]. Alternatively, pathology-associated miRNA concentration changes may coincide with opposite changes in cellular miRNA secretion, thus counterbalancing the effect of altered miRNA expression [30]. Such a correlation is actually observed for one of the best established AD biomarkers; amyloid-beta 42 (Aβ_42_) levels are found to be increased in the brains of individuals with AD, but not in their CSF or blood, where Aβ_42_ shows a marked decrease. Only future work will help to clarify the potential pathogenic and prognostic role of these and other miRNAs highlighted in our study.

Despite the systematic and comprehensive nature of our approach, our study and its results are subject to a range of limitations and potential sources of bias. First, our methods to identify relevant papers and the data reported therein may be erroneous, e.g. owing to imperfect search strategies or mistakes made during the data entry and/or double-checking process. However, by using an adapted version of a carefully crafted data handling protocol already successfully applied to thousands of genetic association studies for different phenotypes [8, 9], we do not expect data entry errors to have largely affected the main conclusions of our study. A related limitation relates to the “quality” of the primary papers which we did not specifically assess here. Poor quality studies (e.g. those applying inadequate data normalisation or analysis strategies) may have affected the meta-analysis outcome to some degree.

The second and likely most important limitation of our study relates to the fact that meta-analyses in this report were based on synthesizing reported *p*-values as test statistic and direction of differential expression (“up” or “down” in AD cases vs controls). This method of quantitative data aggregation may appear relatively crude and simplistic as no direct effect estimates (e.g. fold change) or measures of precision (e.g. standard errors), which are typically utilised in quantitative research syntheses, were used. However, we note that these latter metrics were reported inconsistently and infrequently across the studies included in our analyses precluding effect-size based analyses (and thereby assessments of effect-size heterogeneity across study results) or potential biases, e.g. publication bias or selective reporting bias. In a sense, our analysis method represents the lowest common denominator allowing to aggregate the largest number of datasets per meta-analysis. Only future work adhering to standardised experimental and reporting standards will improve this situation in research aimed at synthesizing gene expression data across different datasets. Until then, the miRNAs highlighted in the meta-analyses presented here likely represent the most promising candidates to show genuine differential expression in AD when compared to controls.

Third, we note that the sample sizes of individual studies aggregated in this work are comparatively small. This is likely due to the fact that the research probing for differential gene expression is still in an early phase with more powerful applications (e.g. mRNA or miRNA sequencing using next-generation technologies) only slowly becoming affordable to be performed on a larger scale. To a degree, the bias inherent in analysing small sample sizes was attenuated here by combining the results of multiple independent small datasets by metaanalysis. However, even our largest analysis (i.e. that of hsa-miR-29a-3p in blood with a combined sample size of 1043 from ten independent studies [Supplementary Table 2]) is small compared to other fields, such as GWAS of complex traits where recent studies aggregated more than one million individuals by meta-analysis [31]. Until much larger miRNA expression studies are published (and meta-analysed), the miRNAs highlighted here likely represent the most promising findings in the field.

Fourth, similar to most other epigenetic mechanisms, evidence of differential expression cannot serve as indicator regarding cause or effect of the underlying pathophysiological process, disease progression, and/or treatment effects. Taking the AD brain as example, it is well established that the cell composition changes as the disease progresses; neuronal cells are depleted while the number of other cell types, e.g. glial cells, can increase. Such changes can give rise to differential miRNA expression, but do not necessarily reflect changes at the cellular level and/or imply the involvement of specific biochemical pathways, which is of primary interest in order to gain further insights into the pathophysiology of a given disease. Future studies could overcome this potential source of bias by measuring expression in specific cells rather than a heterogeneous tissue specimen and by cell selection techniques such as laser capture micro-dissection. Furthermore, research on gene (including miRNA) expression should consistently provide detailed information about disease duration, severity and treatment prior to tissue donation, as this would enable assessment of how these factors may influence brain miRNA expression.

In conclusion, in our systematic and quantitative assessment of more than 100 individual studies we identified 57 unique miRNAs showing study-wide differential expression in AD vs controls. Nearly a quarter of these miRNAs can be considered novel findings as they were not highlighted in either the abstracts of included studies or in recently published qualitative review articles, emphasising the utility of the meta-analytic approach taken here. Taken together, the findings of this work have the potential to substantially advance research on miRNA (dys)function in AD and help pave the way, in concert with non-miRNA markers, for the development of improved pre-disease biomarker panels allowing an early detection and monitoring of the pathophysiologic changes underlying this devastating disease.

## ACKNOWLEDGEMENTS

Author contributions: *Study concept and design:* Takousis, Perneczky, Bertram. *Analysis and interpretation of data:* Takousis, Sadlon, Perneczky, Bertram. *Drafting of the manuscript:* Takousis, Sadlon, Perneczky, Bertram. *Critical revision of the manuscript for important intellectual content:* Takousis, Sadlon, Schulz, Wohlers, Dobricic, Middleton, Lill, Perneczky, Bertram. *Statistical analysis:* Takousis, Sadlon, Schulz, Wohlers, Dobricic, Perneczky, Bertram. *Administrative, technical, and material support:* Schulz, Wohlers, Dobricic, Middleton. *Study supervision:* Perneczky, Bertram.

## Funding sources

This work was supported by the EU (grant number IMI_GA_EMIF-115372 to L.B.), a scholarship of the Peter und Traudl Engelhorn Foundation (to I.W.), an “Exzellenzmedizin” thesis research scholarship of the University of Lübeck (to J.S.), Stevenage Bioscience Catalyst/Imperial Innovations (grant number 7164/SBC014RP to R.P.), and an Imperial College President’s PhD scholarship (to A.S.). All funders of the study had no role in study design, data collection, data analysis, data interpretation, or writing of the report. All authors had full access to all the data in the study and had final responsibility for the decision to submit for publication.

## Declaration of interests

We declare no competing interests.

